# Award or Reward? Which comes first, NIH funding or research impact?

**DOI:** 10.1101/193755

**Authors:** Lucas C. Parra, Lukas Hirsch

## Abstract

Which comes first, funding or research impact? We use causal inference to determine the direction of influence in a record of 70,000 NIH-funded investigators, which was recently released. Contrary to the basic premise of many funding policies, we find that the number of citations of an investigator determine funding levels, but not the other way around.

---

Common sense suggests that increasing research funding increases research productivity. We also know that with increased productivity, the odds of receiving funding increases. So, which comes first, productivity or funding? To answer this question we used causal inference (*1,2*) to analyze data that was recently released (*3*) on over 70,000 principal investigators who had received NIH funding between 1996 and 2014. We find that research productivity, as measured by citation numbers, predicts the level of NIH funding. However, the reverse is not true. There appears to be no simple monotonic relationship between the level of funding and the distribution of research productivity (Fig. 1). As measures of research productivity and funding level, we used metrics previously agreed upon by an NIH working group (*3*). These are the relative citation ratio (RCR), which measures the number of yearly citations of an investigator relative to the number of citations by their peers in the same research field, and the grant support index (GSI), which adds up NIH grants based on a point system. Given the importance of research funding and productivity, it is no surprise that these and other metrics are vigorously debated. We do not endorse, nor question, the merits of these metrics, but simply analyze them to determine which of two alternative causations is better supported by these data. Specifically, productivity determines funding (Hypothesis 1: RCR->GSI), or funding determines productivity (Hypothesis 2: GSI->RCR). The joint distribution of funding and productivity across investigators is shown in Fig. 1A. Causal inference applies Occam’s razor to such observational data as follows: If a given value of variable A straightforwardly determines the distribution of variable B, but there is no simple model for the distribution of variable A given B, then the hypothesis that A causes B is favored, over the hypothesis that B causes A (*1,2*). To explore hypothesis 1 with an established causal inference method (*2*), consider fig. 1B, which shows the mean of GSI conditioned on RCR. Evidently, there is a smooth monotonic increase in average NIH research funding (GSI) as research productivity (RCR) increases. The inverse is not true. As research funding increases, citations averaged across investigators do not follow a smooth monotonic increase (Fig. 1E). When we look at the spread around these mean values, we see that they also increase (Fig. 1C & F). That is, with increasing success, uncertainty also increases. If the spread we observe is due to chance, e.g. the result of a lucky or unlucky year, then we expect that the standard deviation is tightly linked to and monotonically increases with the mean (which is a characteristic of many stochastic processes; see Methods). This is precisely what we observe for funding levels (Fig. 1D) but not for research productivity (Fig. 1G). To summarize, there is a simple model for the hypothesis that citations determine funding, but no simple model to explain the behavior of citations as a function of funding (for a formal hypothesis test see Methods). A similar relationship is found when considering other measures of citations and funding provided by Lauer et al. (2017) (*3*) (see Methods). We conclude that research 1 Send correspondence to LCP. productivity determines NIH funding, but not the other way around. In short, investigators are rewarded rather than awarded by the NIH. Whether this is true for other funding mechanisms or other metrics of research productivity remains to be seen.

**Fig. 1:**
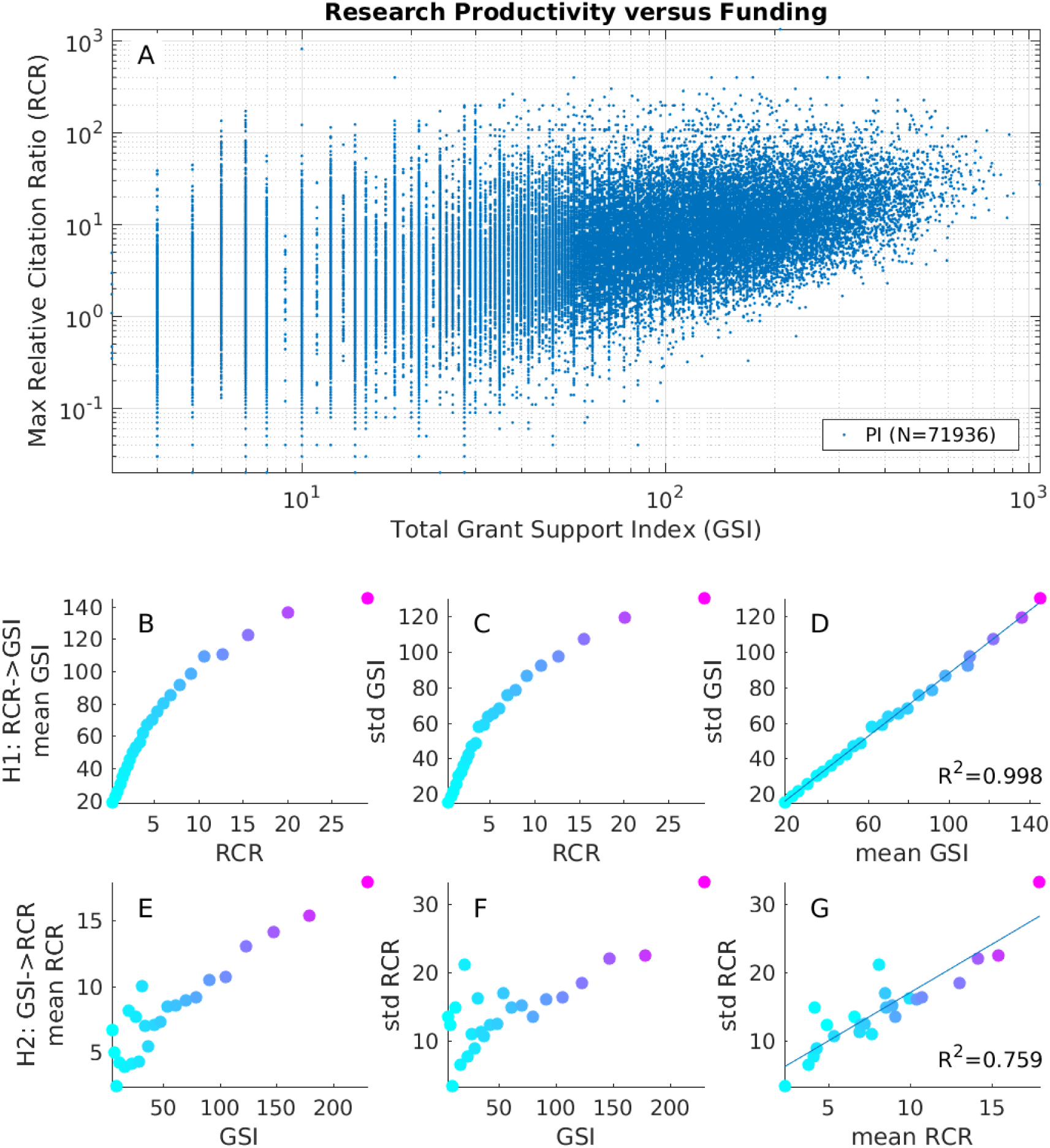
Research productivity versus funding. (A) Distribution of research productivity and funding across NIH funded principal investigators (*3*). Productivity is measured in terms of relative citation ratio (RCR), here as the maximum value over the funding period. Funding is measured in terms of the grant support index (GSI), here as the sum over the funding period. (B & C) Mean and standard deviation of funding (GSI) conditioned on productivity (RCR, binned in 4-percentile intervals, color represents interval; the display excludes the last interval for better visualization). (D) These means and standard deviations of GSI conditioned on RCR follow a linear trend (R^2^ indicates goodness of fit). The smooth monotonic relationship suggests that RCR->GSI is a simple relationship. (E-G) The mean and standard deviation of productivity (RCR) conditioned on funding (GSI) are more irregular, and would therefore require a more complex model to explain GSI->RCR. Code to generate this figure is available here.

NIH funding is disproportionately awarded to senior investigators, and a number of policies aim to address this, such as a relaxed threshold for funding junior investigators, or funding mechanisms that do not require preliminary data. Along these lines, NIH recently considered a rule that would limit funding above a GSI of 21 points. This has caused considerable debate (*4,5*), and the present data was released, in part to argue that productivity does not scale with increasing funding levels (*3*). We were not concerned with the question of how productive researchers are above a GSI of 21, as this affects only a 1.3% of all investigators in these data. Instead, we found a phenomenon that appears to affects the bulk of NIH investigators, namely, that the funding process primarily serves to reward citations. While our finding may be limited to NIH funding and citation records, we suspect that the creativity which inspires citations is not primarily dependent on funding. Instead, other factors such as training, environment, or talent may play a bigger role. It may be that creativity cannot be purchased, although surely some support is necessary to give researchers the time and space necessary to be creative.

## Methods

### Other metrics

Other measures of productivity and funding that were released by Lauer et al. (2017) (*3*) yield similar results. For example maximal RCR vs annual GSI yields R^2^=0.97 for H1, and R^2^=0.87 for H2 (Fig. 2 D&H). Indeed, R^2^ is higher for H1 than for H2 for 28 of 35 possible pairings of the productivity and funding measures. There has been some critique in particular of the way citations are attributed to investigators for NIH program grants (P grants).^6^ When we limit the analysis to recipients of R grants only (e.g. R01 and R21 grants which minimize^7^ the problems noted) we obtain similar result (R^2^ is higher for H1 than for H2 for 29 of 35 possible pairings, for this sample of N=65574 investigators).

**Fig. 2:**
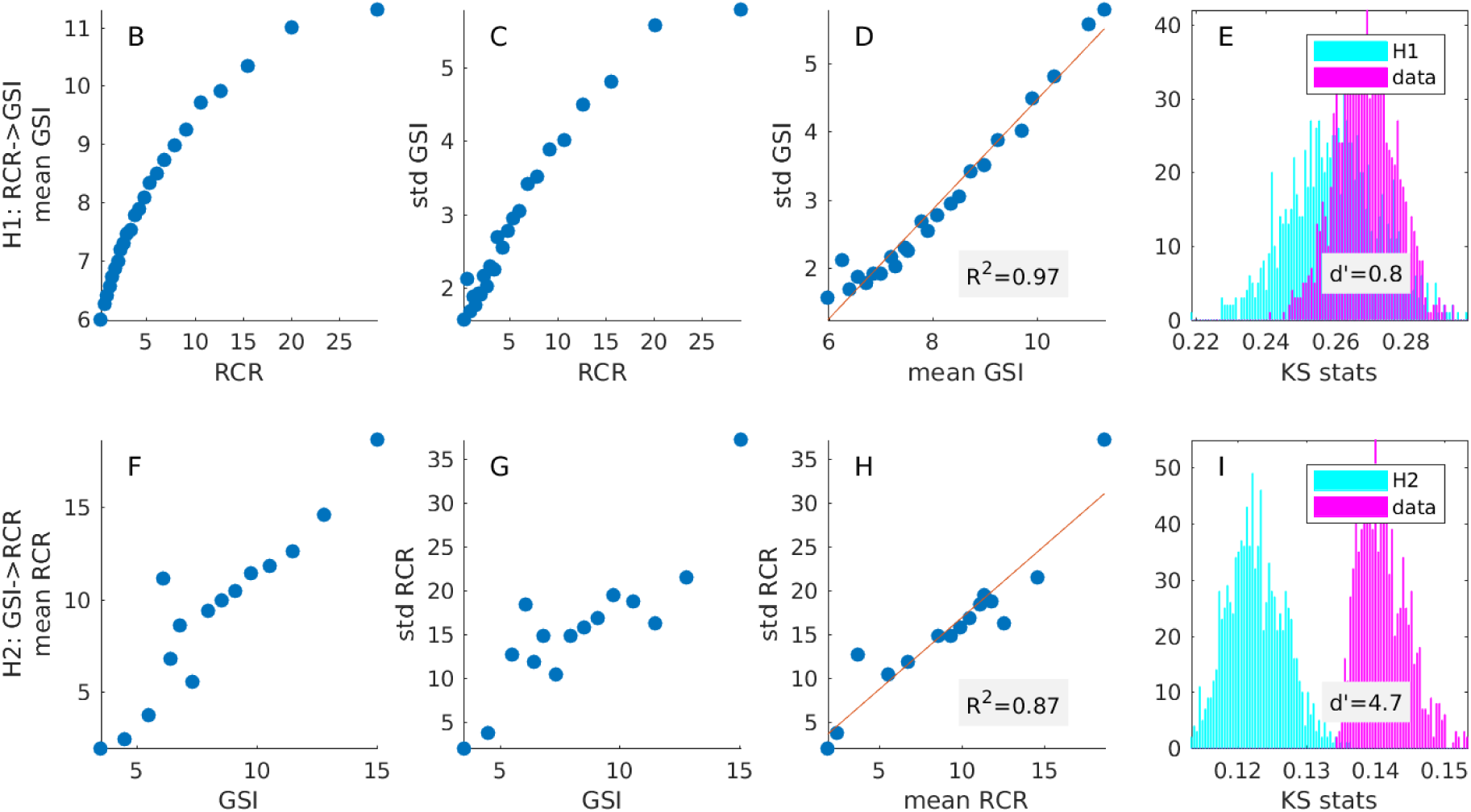
Statistical test for consistency of hypothesized models with the data. Panels A-B and F-G are the same as Fig. 1 but with GSI given as annual values, which is the metric proposed by Lauer et al. (2017) (*3*). New panels E & I show the distribution of KS statistic values obtained for the real data (magenta) vs the model (cyan) with mean and standard deviation adjusted according to the linear model in panels D and H. KS statistics measure how different the two conditional distributions are across different intervals of the conditioning variable, averaged across intervals. The comparison of the conditional samples picks a reference interval at random, and this random selection is repeated 10^5^ times, hence there is a range of values for the KS statistics. The separation of these KS values is measured as d-prime. High d-prime indicates that the data differs from the hypothesized distributions. Code to generate this figure is available here.

### Hypothesis test

The simple model proposed by this analysis is that as citation count increases, the mean chance of obtaining funding increases in a smooth and orderly fashion, yet the actual number of grants is otherwise left to chance. Chance distributions that are determined by their mean, with a monotonic increasing deviation, are characteristic of a number of stochastic processes such as Poisson, geometric, or lognormal. In fact, the linear trend in Fig. 1D is suggestive of a lognormal process. (Gaussian noise with constant deviation, as often assumed in causal inference,^1^ is not applicable here.) For the statistical test we will assume that conditional distributions are independent of the conditioning variable, except for a changing mean and a linearly dependent standard deviation. For H1 the conditioning variable is the number of citations, and the conditioned variable is funding (Fig. 2B-D); for H2 it is the reverse (Fig. 2F-H). If the model is correct, then all intervals of the conditioning variable (centered at points in panels B-D or F-H) should have comparable distributions for the conditioned variable (after adjusting mean and deviation). We measure the difference of conditional samples between different intervals of the conditioning variable using the Kolmogorov-Smirnov (KS) statistic. By selecting intervals at random while adjusting the mean and deviation according to the linear model we establish the chance distribution of KS values (Fig. 2E or I), under the hypothesized model (cyan), and for the real data (magenta). The two distributions are overlapping for H1 (d-prime=0.8) suggesting that the data is consistent with the corresponding model. However, they do not overlap for H2 (d-prime=4.7) indicating that the model does not capture the data well (see <u>code</u> for detail). In total, this metric favors H1 over H2 in 31 of 35 of possible pairings of the production versus funding variables (and 30 of 35 pairings when restricting analysis for R grants).

## References

1. P. O. Hoyer, D. Janzing, J. M. Mooij,J. Peters, B. Schölkopf, Nonlinear causal discovery with additive noise models. Advances in Neural Information Processing Systems 21, 689–696 (2009).

2. L. Muchnik, et al, “Origins Of Power-Law Degree Distribution In The Heterogeneity Of Human Activity In Social Networks”. Scientific Reports 3.1 (2013).

3. M. Lauer,D. Roychowdhury, K. Patel, R. Walsh,K. Pearson, Marginal Returns And Levels Of Research Grant Support Among Scientists Supported By The National Institutes Of Health (2017).

4. J. Kaiser, NIH abandons controversial plan to cap grants to big labs, creates new fund for younger scientists. Science (2017). doi:10.1126/science.aan6947

5. S. Reardon, NIH grant limits rile biomedical research community. Nature 545, 142–143 (2017).

6. S. Crotty, Considering assessments of scientific productivity and ‘ghost authors’, Medium, Jun 8, 2017, URL

7. K. W. Boyack, P. Jordan, Metrics associated with NIH funding: a high-level view, J Am Med Inform Assoc, 18: 423e431 (2011).

